# SARS-COV-2 Delta variant displays moderate resistance to neutralizing antibodies and spike protein properties of higher soluble ACE2 sensitivity, enhanced cleavage and fusogenic activity

**DOI:** 10.1101/2021.11.05.467523

**Authors:** Sabari Nath Neerukonda, Russell Vassell, Sabrina Lusvarghi, Richard Wang, Fernando Echegaray, Lisa Bentley, Ann E. Eakin, Karl J. Erlandson, Leah C. Katzelnick, Carol D. Weiss, Wei Wang

## Abstract

The SARS-CoV-2 B.1.617 lineage variants, Kappa (B.1.617.1) and Delta (B.1.617.2, AY) emerged during the second wave of infections in India, but the Delta variants have become dominant worldwide and continue to evolve. The spike proteins of B.1.617.1, B.1.617.2, and AY.1 variants have several substitutions in the receptor binding domain (RBD), including L452R+E484Q, L452R+T478K, and K417N+L452R+T478K, respectively, that could potentially reduce effectiveness of therapeutic antibodies and current vaccines. Here we compared B.1.617 variants, and their single and double RBD substitutions for resistance to neutralization by convalescent sera, mRNA vaccine-elicited sera, and therapeutic neutralizing antibodies using a pseudovirus neutralization assay. Pseudoviruses with the B.1.617.1, B.1.617.2, and AY.1 spike showed a modest 1.5 to 4.4-fold reduction in neutralization titer by convalescent sera and vaccine-elicited sera. In comparison, similar modest reductions were also observed for pseudoviruses with C.37, P.1, R.1, and B.1.526 spikes, but seven- and sixteen-fold reduction for vaccine-elicited and convalescent sera, respectively, was seen for pseudoviruses with the B.1.351 spike. Four of twenty-three therapeutic neutralizing antibodies showed either complete or partial loss of neutralization against B.1.617.2 pseudoviruses due to the L452R substitution, whereas six of twenty-three therapeutic neutralizing antibodies showed either complete or partial loss of neutralization against B.1.617.1 pseudoviruses due to either the E484Q or L452R substitution. Against AY.1 pseudoviruses, the L452R and K417N substitutions accounted for the loss of neutralization by four antibodies and one antibody, respectively, whereas one antibody lost potency that could not be fully accounted for by a single RBD substitution. The modest resistance of B.1.617 variants to vaccine-elicited sera suggest that current mRNA-based vaccines will likely remain effective in protecting against B.1.617 variants, but the therapeutic antibodies need to be carefully selected based on their resistance profiles. Finally, the spike proteins of B.1.617 variants are more efficiently cleaved due to the P681R substitution, and the spike of Delta variants exhibited greater sensitivity to soluble ACE2 neutralization, as well as fusogenic activity, which may contribute to enhanced spread of Delta variants.

## 1. Introduction

Since its origin in December 2019, severe acute respiratory syndrome coronavirus 2 (SARS-CoV-2) has spread globally to cause a coronavirus disease 2019 (COVID-19) pandemic that recorded more than 248 million infections and claimed 5.0 million lives thus far (Johns Hopkins Coronavirus Resource Center; https://coronavirus.jhu.edu). SARS-CoV-2 trimeric spike (S) glycoprotein on the virion surface binds the angiotenisin converting enzyme (ACE2) to facilitate cellular entry and is the target of therapeutic neutralizing antibodies (nAbs) and vaccines [1–6]. The spike is proteolytically processed between R^685^ | S^686^ into S1 and S2 subunits, which facilitates subsequent cleavage at the S2’ site (R^815^ | S^816^) by TMPRSS2 for viral entry into respiratory cells. The S1 subunit spans the N-terminal domain (NTD) and the receptor binding domain (RBD) within the C-terminal domain (CTD) whereas the S2 subunit spans the fusion peptide and a linker region flanked by heptad repeat regions that drive virus-cell membrane fusion [7]. Besides virus-cell fusion, spike protein on the cell surface can trigger receptor-dependent syncytia formation via cell-cell fusion.

Currently available vaccines and therapeutic antibodies target the spike glycoprotein of an early isolate of SARS-CoV-2. The continued evolution of SARS-CoV-2 resulted in the emergence of several variants of distinct lineages globally, raising concerns over variant transmissibility and immune escape. One of the earliest variants that is highly infectious and thus, became globally dominant is B.1 (D614G) [8–11]. However, convalescent sera from individuals infected with an early viral isolate (Wuhan-Hu-1) effectively cross neutralized D614G [12]. Subsequent genomic surveillance has led to the identification of several convergently evolving lineages, including in UK-B.1.1.7 (Alpha), South Africa-B.1.351 (Beta), Brazil/Japan-P.1 (Gamma), California USA-B.1.427/B.1.429 (Epsilon), Northeast USA-B.1.526 (Iota), USA/Japan (R.1), Peru/Chile-C.37 (Lambda), and Liverpool-A23.1. By late 2020, B.1.617 lineage variants emerged in India and have spread rapidly throughout the world. The Kappa (B.1.617.1) variant emerged early in the second wave, followed by the Delta (B.1.617.2) and its sub-lineage (AY.1 and AY.2) variants, which are currently dominant in many parts of the world.

Several key substitutions in the RBD of spike were demonstrated to either enhance affinity towards ACE2 or contribute to immune escape. The E484K substitution in the RBD of B.1.351, P.1, R.1, and B.1.526 variants was previously identified among *in vitro* escape mutants selected against single antibody and antibody cocktails [13,14]. Several studies have described a significant drop in the neutralization potency of convalescent and vaccine sera, as well as numerous therapeutic nAbs that contacted the mutated sites in B.1.351, P.1, and B.1.526 lineages, particularly E484K [15–17]. The B.1.427/B.1.429 and B.1.617 lineage variants share the L452R substitution in RBD [18]. The L452R substitution has been demonstrated to enhance ACE2 binding and pseudovirus infectivity [19] and reduce or ablate the neutralizing potency of 10 out of 34 RBD-specific mAbs tested [18]. The E484Q along with L452R is present in the RBD of B.1.617 sub-lineages, B.1.617.1 and B.1.617.3. B.1.617.1 contains additional substitutions in the NTD (T95I), the NTD antigenic supersite β-hairpin (G142D and E154K), within the S1/S2 cleavage junction (P681R), and in the S2 subunit (Q1071H) [20]. E484Q was also previously identified as an RBD escape mutant for an RBD-specific mAb [21,22].

The spike protein of the B.1.617.2 variant contains nine substitutions and deletions compared to the early D614G variant used here as wild type (WT or D614G). The three substitutions (T19R, G142D, and R158G) and two deletions (ΔE156, ΔF157) in NTD occur in the NTD antigenic supersite spanning between residues 14–20, 140–158 and 245–264 [23]. In addition, the B.1.617.2 spike also has two substitutions in the receptor binding domain (RBD) (L452R, T478K), one substitution proximal to S1/S2 cleavage site (P681R), and one in the S2 region (D950N). An additional substitution in RBD, K417N is also observed in the RBD of AY.1. The L452R RBD substitution was shown to confer modest resistance to neutralization by convalescent sera, vaccine-elicited sera, and therapeutic neutralizing anti-bodies in the context of other variants, such as B.1.427/B.1.429, and B.1.617.1 [19,24]. The E484K substitution found in B.1.351, B.1.526, P.1, P.3 (theta) and some B.1.617 variants, confers some level of resistance to neutralization by convalescent sera, vaccine-elicited sera, and therapeutic neutralizing antibodies [15]. In addition to neutralizing antibody resistance, spike mutations affect proteolytic processing, virion incorporation, ACE2 affinity, as well as membrane fusion [25–27].

The global dominance and ongoing evolution of B.1.617 lineage variants require continuing assessment of the neutralization potency of convalescent sera, vaccine-elicited sera, and therapeutic neutralizing antibodies against emerging B.1.617 variants. Here, we measured the neutralization potency of convalescent sera, vaccine-elicited sera, and therapeutic neutralizing antibodies against two independent variants each in the Kappa (B.1.617.1) and Delta (B.1.617.2, AY.1) lineages and assessed the contribution of the RBD substitutions in conferring resistance. We found that resistance to convalescent and vaccine-elicited sera was predominantly conferred by RBD substitutions E484Q, T478K, and L452R. Furthermore, out of 23 therapeutic neutralizing antibodies tested, Kappa and Delta pseudoviruses displayed complete resistance to 5 neutralizing antibodies and partial resistance to one antibody. Finally, we show that the P681R substitution confers enhanced furin processing in spike protein of B.1.617 lineage variants that corresponded to enhanced cell-cell fusion activity. However, only Delta spike protein exhibited greater sensitivity to soluble ACE2 inhibition, implying enhanced ACE2 affinity. This feature along with enhanced cell-cell fusion activity may contribute to the dominance of the B.1.617.2 variant.

## 2. Materials and Methods

### 2.1. Ethics Statement

Use of de-identified sera and nAbs in this study was approved by the U.S. Food and Drug Administration Research in Human Subjects Committee. Vaccine-elicited sera were collected at the U.S. Food and Drug Administration with written consent under an approved Institutional Review Board (IRB) protocol (FDA IRB Study # 2021-CBER-045).

### 2.2. Plasmids and cell lines

Codon-optimized full-length open reading frames of the S genes of SARS-COV-2 variants were cloned into pcDNA3.1(+) or pVRC8400 by GenScript (Piscataway, NJ). The spikes of variants used in this study were listed in Table 1. The HIV gag/pol (pCMVΔR8.2), and Luciferase reporter (pHR’CMV-Luc) plasmids were obtained from the Vaccine Research Center (National Institutes of Health, Bethesda, MD) [28,29]. 293T-ACE2.TMPRSS2 cells stably expressing human angiotensin converting enzyme 2 (ACE2) and transmembrane serine protease 2 (TMPRSS2) were previously described (BEI Resources Cat no: NR-55293) [30]. The 293T and 293T-ACE2.TMPRSS2 cells were maintained at 37°C in Dulbecco’s modified eagle medium (DMEM) supplemented with high glucose, L-Glutamine, minimal essential media (MEM) non-essential amino acids, penicillin/streptomycin and 10% fetal bovine serum (FBS).

**Table 1.** List of spike variants used in the present study

| Variants | Name used in this report | Amino acid substitutions in spike compared to Wuhan-Hu-1 | RBD substitutions |
| --- | --- | --- | --- |
| Wuhan-Hu-1 | Wuhan-Hu-1 |  |  |
| Wuhan-Hu-1-PP681R | Wuhan-Hu-1+P681R | P681R |  |
| B.1 | WT(D614G) | D614G |  |
| B.1-P681R | P681R | D614G, P681R |  |
| B.1-P681H | P681H | D614G, P681H |  |
| B.1-L452R | L452R | L452R, D614G | L452R |
| B.1-E484Q | E484Q | E484Q, D614G | E484Q |
| B.1-T478K | T478K | T478K, D614G | T478K |
| B.1-K417N | K417N | K417N, D614G | K417N |
| B.1-L452R-T478K | L452R+T478K | L452R, T478K, D614G | L452R, T478K |
| B.1-L452Q-F490S | L452Q+F490S | L452Q, F490S, D614G | L452Q, F490S |
| B.1.617.1 (Kappa) | B.1.617.1 (A) | G142D, E154K, V382L, L452R, E484Q, D614G, P681R, Q1071H, D1153Y | V382L, L452R, E484Q |
| B.1.617.1 (Kappa) | B.1.617.1 (B) | T95I, G142D, E154K, L452R, E484Q, D614G, P681R, Q1071H | L452R, E484Q |
| B.1.617.2 (Delta) | B.1.617.2 | T19R, G142D, E156Δ, F157Δ, R158G, L452R, T478K, D614G, P681R, D950N | L452R, T478K |
| AY.1 (Delta plus) | AY.1 | T19R, T95I, G142D, E156Δ, F157Δ, R158G, W258L, K417N, L452R, T478K, D614G, P681R, D950N | K417N, L452R, T478K |
| C.37 (Lambda) | C.37 | G75V, T76I, Δ246-252, D253N, L452Q, F490S, D614G, T859N | L452Q, F490S |
| B.1.429 (Epsilon) | B.1.429 | S13I, P26S, W152C, L452R, D614G | L452R |
| R.1 | R.1 | W152L, E484K, D614G, G769V | E484K |
| B.1.526 (Iota) | B.1.526 | L5F, T95I, D253G, E484K, D614G, A701V | E484K |
| P.1 (Gamma) | P.1 | L18F, T20N, P26S, D138Y, R190S, K417T, E484K, N501Y, D614G, H655Y, T1027I, V1176F | K417T, E484K, N501Y |
| B.1.1.7 (Alpha) | B.1.1.7 | 69-70Δ, Y144Δ, N501Y, A570D, D614G, P681H, T716I, S982A, D1118H | N501Y |
| B.1.351 (Beta) | B.1.351 | L18F, D80A, D215G, 241-243Δ, K417N, E484K, N501Y, D614G, A701V | K417N, E484K, N501Y |

### 2.3. Human sera and therapeutic neutralizing antibodies

Convalescent sera from SARS-COV-2 infected donors (n=10) collected 6-61 days after symptom onset were purchased from Bocabiolistics (Bocabiolistics, FL). Donors were 18-73 years old with six males/four females. The information about the convalescent sera is shown in Table 2. Sera from Pfizer BNT162b2- (n=15) or Moderna mRNA 1273-vaccinated individuals (n=14) obtained two weeks after the second vaccination were used in this study. Vaccinated individual donors were 21-65 years old with six males/nine females for Pfizer BNT162b2 vaccination and 8 males/6 females for Moderna vaccination. All sera were tested negative for non-specific neutralization using amphotropic murine leukemia enveloped pseudovirus. Vaccinated donors were prescreened for absence of both history of SARS-COV-2 infection and SARS-CoV-2 neutralizing antibodies prior to vaccination. Twenty-three therapeutic neutralizing antibodies against SARS-COV-2 spike protein were generously donated by different pharmaceutical companies for the U.S. Government COVID-19 response therapeutics research team efforts to define neutralization profiles against existing and emerging SARS-CoV-2 variants [17]. Due to a confidentiality agreement with the manufacturers, neutralizing antibodies described are shown with blinded identification codes as follows: single neutralizing antibodies (nAbs A to R), combination of two neutralizing antibodies (cnAbs S to X), and polyclonal neutralizing antibodies (pnAbs III to IV).

**Table 2.** Demographics and infection history of convalescent sera donor individuals

| Donor ID | Gender | Age | Days from 1<sup>st</sup> symptoms | Residue substitutions in spike of SARS-CoV-2-infected individual |
| --- | --- | --- | --- | --- |
| 1 | Male | 36 | 16 | D614G |
| 2 | Female | 29 | 28 | D614G |
| 3 | Male | 70 | 14 | D614G |
| 4 | Female | 54 | 61 | D614G |
| 5 | Male | 18 | 13 | D614G |
| 6 | Male | 59 | 9 | D614G |
| 7 | Male | 25 | 40 | S13I, Q52R, A67V, 69-70del, 144del, L452R |
| 8 | Female | 73 | 59 | L452R, D614G |
| 9 | Female | 33 | 26 | L452R, D614G |
| 10 | Male | 55 | 6 | W152C, L452R, D614G |

### 2.4. soluble ACE2 production

The production of His-tagged soluble ACE2 was performed in FreeStyle^™^ 293-F cells and purified using HisPur^™^ Ni-NTA Resin (Thermo Scientific) as described previously [31].

### 2.5. SARS-CoV-2 pseudovirus production and neutralization assay

HIV-based lentiviral pseudoviruses with spike proteins were generated as previously described [30]. The B.1 spike containing D614G, was used as wild type (WT(D614G)). Briefly, pseudoviruses bearing the spike glycoprotein and packaging a firefly luciferase (FLuc) reporter gene were produced in 293T cells by co-transfection of 5μg of pCMVΔR8.2, 5μg of pHR’CMVLuc and 4μg of pcDNA3.1(+) or 0.5 ug of pVRC8400-encoding a codon-optimized spike gene with the desired substitutions. Pseudovirus supernatants were collected approximately 48 hr post-transfection, filtered through a 0.45μm low protein binding filter, and stored at −80°C. Neutralization assays were performed using 293T-ACE2.TMPRSS2 cells in 96 well plates as previously described [30]. Briefly, pseudoviruses with titers of approximately 1×10^6^ relative luminescence units (RLU)/ml of luciferase activity were incubated with serially diluted sera or antibodies for two hr at 37°C. Pseudovirus and serum or antibody mixtures (100 μl) were then inoculated onto the plates pre-seeded one day earlier with 3.0 x 10^4^ cells/well. Pseudovirus infectivity was scored 48 hr later for luciferase activity. The antibody concentration or inverse of the sera dilutions causing a 50% reduction of RLU compared to pseudovirus control were reported as the neutralization titer. Titers were calculated using a nonlinear regression curve fit (GraphPad Prism software Inc., La Jolla, CA). The mean titer from at least two independent experiments each with intra-assay duplicates was reported as the final titer. WT(D614G) pseudovirus was run as a control for every assay.

For ACE2 neutralization assay, serially diluted recombinant human soluble ACE2 was incubated with indicated pseudovirus (~1×10^6^ RLU / ml) for one hour at 37°C and 100 μl of pseudovirus and soluble ACE2 mixture was added to 293T-ACE2.TMPRSS2 cells. Luciferase activity was measured 48 hr later. The soluble ACE2 concentration causing a 50% reduction of RLU compared to pseudovirus control were reported as the 50% inhibitory concentration or IC_50_.

### 2.6. Western Blotting

1.25 ml of pseudoviruses were pelleted at 4°C for 2 hr at 15000 rpm using Tomy TX-160 ultracentrifuge. Pseudovirus pellet was resuspended in 1X Laemmli loading buffer and heated at 70°C for 10 minutes. Samples were resolved by 4-20% SDS PAGE and transferred onto nitrocellulose membranes. Membranes were probed for SARS-CoV-2 S1 using rabbit polyclonal antibodies against the RBD domain (Sino Biological; Cat: 40592-T62).

### 2.7. Cell-Cell fusion assay

For measuring spike protein-mediated cell-cell fusion, β-Gal complementation assay was performed as described previously [32]. Briefly, β-Gal ω subunit-expressing 293T cells were transfected with 1μg spike plasmids, whereas 293T-ACE2.TMPRSS2 cells were transfected with β-Gal α subunit using the Fugene 6 reagent. At 24 hr post transfection, the transfected cells were detached using a nonenzymatic cell dissociation solution (Sigma) and washed with DMEM. Spike-transfected/β-Gal ω subunit-expressing 293T cells were mixed with β-Gal α subunit-transfected/293T-ACE2.TMPRSS2 cells at 1:1 ratio to a total of 6 x 10^4^ cells per well on a 96-well plate. The cells were co-cultivated for 24 hr at 37°C. The culture supernatants were then removed, and cell-cell fusion was scored by determination of the β-Gal activity in cocultured cell lysates using a Galacto-Star kit (Applied Biosystems) according to the manufacturer’s instructions.

To ensure equivalent amount of spike cell surface expression levels among treatments, spike-transfected β-Gal ω subunit-expressing 293T cells were quantified for cell surface spike levels by flow cytometry. Spike-transfected 293T cells employed in cell-cell fusion assay were concurrently stained with SARS-CoV-2 positive human polyclonal sera at 1:20 dilution, washed twice and then incubated with FITC-conjugated goat anti-human (KPL Inc., Gaithersburg, MD). The cells were washed twice and then fixed with 2% para-formaldehyde. The results were acquired using BD LSRFortessa^™^ X-20 Cell Analyzer (BD biosciences). The mean fluorescence intensities of spike positive cells were recorded.

### 2.8. Antigenic Cartography

We created a geometric interpretation of neutralization titers against the tested SARS-CoV-2 pseudo-viruses using Racmacs antigenic cartography software (https://github.com/acorg/Racmacs) [33,34]. The map is presented on a grid in which each square indicates one antigenic unit, corresponding to a two-fold dilution of the antibody in the neutralization assay. Antigenic distance is measured in any direction on the map.

### 2.9. Furin prediction score calculations

The prediction of furin-specific cleavage site in spike proteins were computed using the ProP 1.0 Server hosted at http://www.cbs.dtu.dk/services/ProP/ and the PiTou V3 software hosted at http://www.nuo-lan.net/reference.html.

### 2.10. Statistics analysis

One-way analysis of variance (ANOVA) with Dunnett’s multiple comparisons tests (variants compared to WT(D614G)), Mann-Whitney test for the comparison of two groups with unmatched pairs (Pfizer BNT162b2 compared to Moderna) and geometric mean titers (GMT) with 95% confidence intervals were performed using GraphPad Prism software. The *p* values of less than 0.05 were considered statistically significant. All neutralization titers were log2 transformed for analyses.

## 3. Results and Discussion

### 3.1. Neutralization of B.1.617 pseudoviruses by convalescent sera

We first investigated the cross-neutralization potency of convalescent sera from individuals infected with SARS-COV-2 in the U.S. against pseudoviruses bearing spikes of B.1.617.1 and B.1.617.2 variants and their corresponding RBD mutations (Figure 1A). Titers against B.1.617.1 (B), AY.1, E484Q and L452R+T478K pseudoviruses were significantly different from the titers against WT(D614G). Compared to titers against WT(D614G) pseudoviruses (GMT 392), titers against B.1.617.1 (B) pseudoviruses were approximately fourfold lower (GMT 90), confirming and extending other reports [20]. Neutralization titers against WT(D614G) and L452R pseudoviruses were comparable (GMT titers 392 and 364, respectively), while neutralization titers against E484Q pseudoviruses were lower (GMT 165). Titers against B.1.617.2 (GMT 259) and AY.1 (GMT 203) pseudoviruses also showed a slight 1.5- and 1.9-fold reduction, respectively, compared to WT(D614G) pseudoviruses. Against pseudoviruses bearing spikes with T478K substitution in RBD, neutralization titers (GMT 270) were also slightly reduced compared to WT (D614G) (GMT 392). A further reduction in neutralization titers was seen against pseudoviruses bearing both L452R and T478 substitutions in RBD displayed (GMT 192) compared to WT(D614G) (GMT 392).

**Figure 1.**
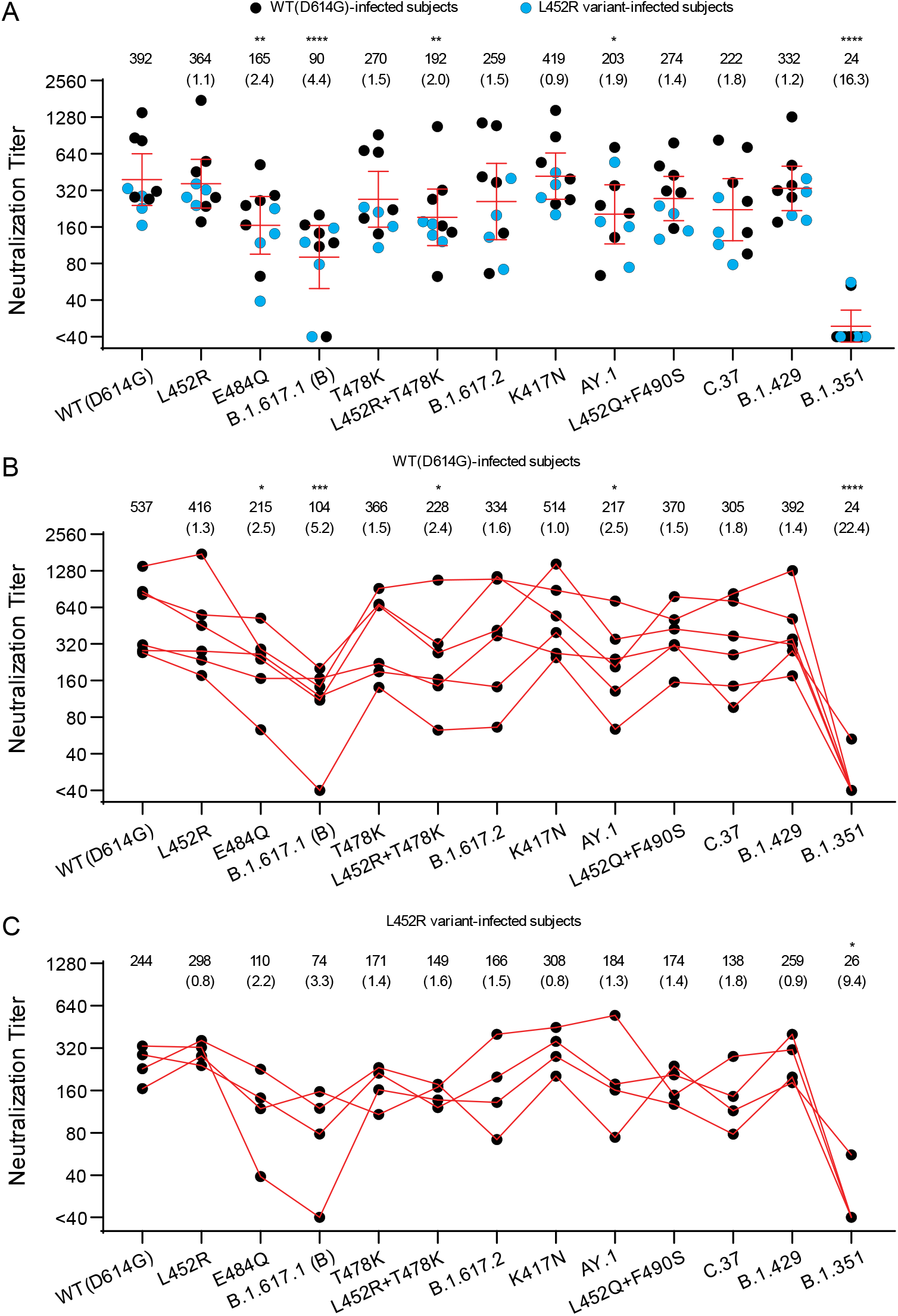
Neutralization of variant SARS-CoV-2 pseudoviruses by convalescent sera. The neutralization titers represented as 50% inhibitory concentrations (IC_50_) against pseudoviruses bearing spike proteins from the indicated variants are plotted. (**A**) Individual neutralization titers of convalescent sera are presented. Blue dots: sera from subjects infected with variants containing L452R in spike. Black dots: sera from subjects infected with WT(D614G) variants. (**B**) The neutralization titers of individuals infected with WT(D614G) SARS-COV-2. (**C**) The neutralization titers of individuals infected with SARS-COV-2 bearing L452R in spike. The numbers over each graph indicate the GMT. The numbers in parentheses are the ratios of WT(D614G) GMT/individual variant GMT. P values were calculated by One-way analysis of variance (ANOVA) with Dunnett’s multiple comparisons tests (variants compared to WT(D614G)). Titers measuring below the lowest serum dilution of 1:40 were treated as 20 for statistical analysis. All neutralization titers were log_2_ transformed before test. Bars: geometric means of titers (GMT) with %95 CI. *: P≤0.05; **: P≤0.01; ***: P≤0.001; ****: P≤0.0001.

The C.37 variant also has a substitution at L452 residue (L452Q instead of L452R) along with F490S in the RBD. A 1.8-fold reduction in titers against C.37 pseudoviruses was observed compared WT(D614G) pseudoviruses (GMT titers 222 and 392, respectively). A similar 1.4-fold reduction in titers was observed for pseudoviruses with only the L452Q and F490S substitutions, indicating that these RBD substitutions may contribute to C.37 resistance. In comparison with other variants, we found that titers against B.1.429 pseudoviruses (GMT 332) were comparable to WT(D614G), but titers against the B.1.351 pseudovirus were 16.3-fold reduced (GMT 24) compared to WT(D614G), consistent with prior reports showing marked reductions of cross-neutralization against this variant [35–37].

Because the convalescent sera came from individuals that previously were infected by different variants (Table 2), we also explored differences in the neutralization titers between those infected by D614G variants lacking L452R and those infected by variants containing L452R (all have L452R and D614G, except one lacking D614G). The small number of samples in each group precludes conclusions, but we noticed modest differences against some variants. In the group infected by D614G variants that lacked L452R, a 5.2-fold, 1.6-fold, and 2.5-fold reduction in cross-neutralization potency was seen against B.1.617.1 (B), B.1.617.2, and AY.1 pseudoviruses, respectively (Figure 1B). The E484Q substitution in B.1.617.1 (B) only partially contributed to escape from neutralization (GMT 215) with a 2.5-fold reduction in neutralization compared to WT(D614G). Pseudoviruses with single L452R and T478K substitutions each conferred a similar fold change in resistance (1.3- and 1.5-fold, respectively) as B.1.617.2 (1.6-fold) compared to WT(D614G), although a 2.4-fold reduction was seen against the pseudoviruses with the dual L452R+T478K substitutions. This latter finding suggests that substitutions outside of the RBD may be modifying resistance to B.1.617.1 (B) in those infected individuals.

Sera from the group infected by variants that had the L452R substitution showed similar fold changes in neutralization titers as the group infected by WT(D614G) variants without the L452R except against B.1.617.1 (B) and AY.1 pseudoviruses. Fold changes in neutralization titers against B.1.617.1 (B) and AY.1 variants were 3.3 and 1.3, respectively, for the L452 group compared to 5.2 and 2.5, respectively, for the WT(D614G) group (Figure 1B and 1C). The fold reduction of neutralization titers against variants containing substitutions at L452 position, including B.1.617.2, AY.1, B.1.617.1, C.37 and L452R RBD substitution mutant, compared to WT(D614G) also trended lower in the group infected by the L452R variant compared to the group infected by the WT(D614G) variant. The generally lower titers and small numbers of samples in the L452R group, however, may impact the fold changes.

### 3.2. Neutralization of B.1.617 variant pseudoviruses by vaccine-elicited sera

We next assessed the neutralization potency of mRNA vaccine-elicited sera against WT(D614G) and B.1.617 variant pseudoviruses. Sera from fourteen individuals who received two doses of Moderna mRNA-1273 vaccine and fifteen individuals who received two doses of Pfizer/BioNtech BNT162b2 vaccine were collected approximately two weeks after the second immunization. Each vaccine-elicited serum had high neutralization titers against WT(D614G) pseudoviruses, ranging between 578-3935 for Pfizer/BioNtech BNT162b2 and 651-5853 for Moderna (Figure 2A and 2B) (Pfizer/BioNtech BNT162b2 vs Moderna, *p* = 0.1225, Mann-Whitney test). The Moderna vaccine-elicited sera trended towards higher neutralization titers against most variants compared to Pfizer/BioNtech BNT162b2, possibly due to higher vaccine mRNA content and greater interval between priming and boosting for Moderna (4 weeks vs 3 weeks for Pfizer/BioNtech BNT162b2) [38]. Compared to WT(D614G), similar reduction in neutralization towards B.1.617.1, B.1.617.2 and AY.1 was noticed for both Pfìzer/BioNtech BNT162b2 and Moderna vaccine-elicited sera. The average neutralization potency of Pfizer/BioNtech BNT162b2 vaccine-elicited sera was 2-2.5-fold lower against B.1.617.1 pseudoviruses (GMT 642 for B.1.617.1 (A); GMT 519 for B.1.617.1 (B)) compared to WT(D614G) (GMT 1310) and 1.9-2.8-fold lower for B.1.617.2 pseudoviruses (GMT 693 for B.1.617.2; GMT 469 for AY.1 pseudovirus) compared to WT(D614G) (GMT1310) (Figure 2A). The average neutralization potency of Moderna vaccine-elicited sera was 2-2.4-fold reduced against B.1.617.1 (GMT 1019 for B.1.617.1 (A); GMT 856 for B.1.617.1 (B)) compared to WT(D614G) (GMT2015) and 1.8-3.4-fold lower against B.1.617.2 pseudoviruses (GMT 1095 for B.1.617.2; GMT 597 for AY.1) (Figure 2B).

**Figure 2.**
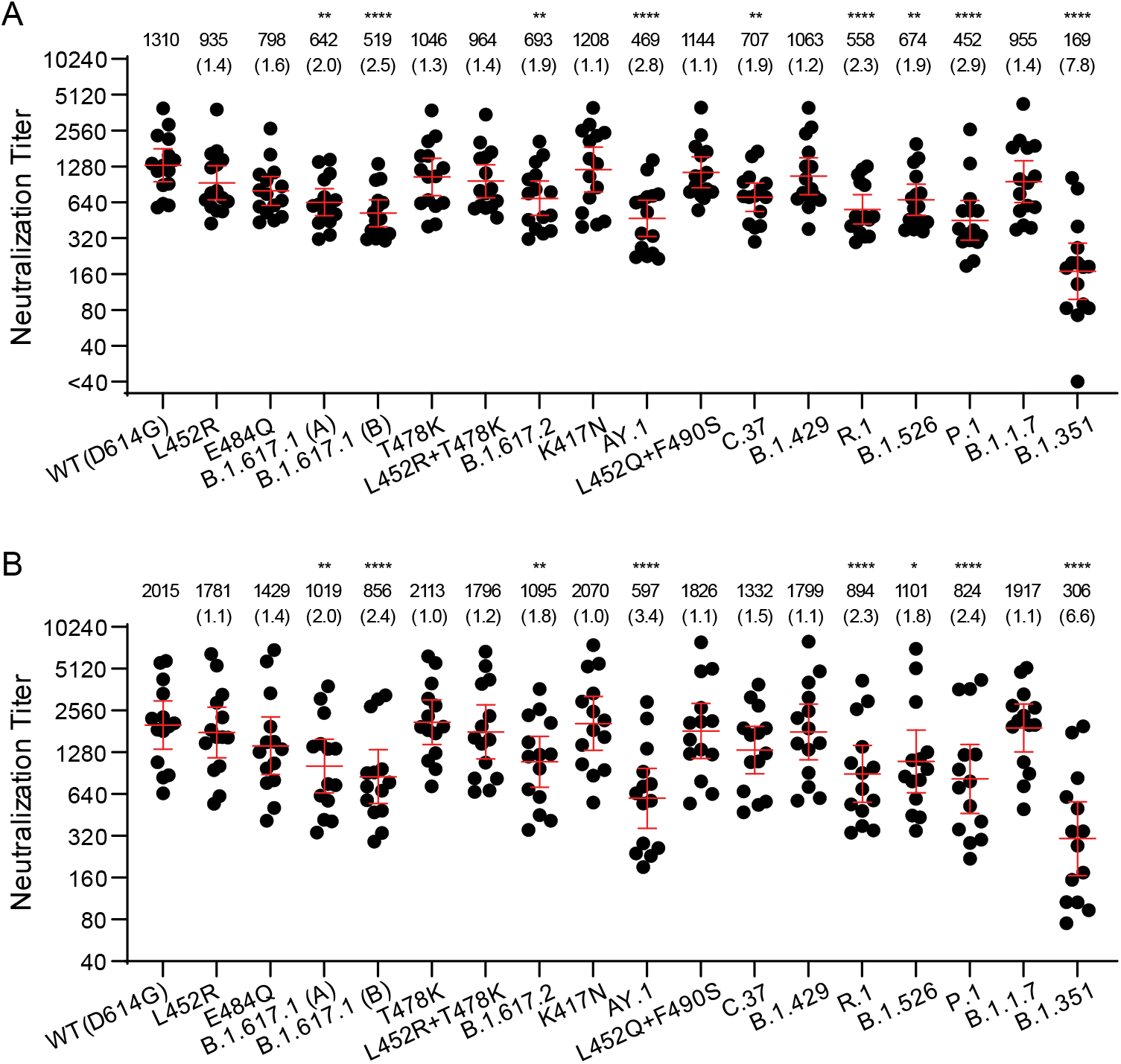
Neutralization of variant SARS-CoV-2 pseudoviruses by vaccine-elicited sera. The neutralization titers represented as 50% inhibitory concentrations (IC_50_) against pseudoviruses bearing spike proteins from the indicated variants are plotted. (**A**) Neutralization titers of Pfizer/BioNtech BNT162b2 vaccination sera are presented. (**B**) Neutralization titers of Moderna mRNA-1273 vaccination sera are presented. Bars: geometric means of titers (GMT) with 95% CI. The numbers over each graph indicate the GMT. The numbers in parentheses are the ratios of WT(D614G) GMT/individual variant GMT. P values were calculated by One-way analysis of variance (ANOVA) with Dunnett’s multiple comparisons tests (variants compared to WT(D614G)). Titers measuring below the lowest serum dilution of 1:40 were treated as 20 for statistical analysis. All neutralization titers were log_2_ transformed before test. *: P≤0.05; **: P≤0.01; ***: P≤0.001; ****: P≤0.0001.

We also investigated the contribution of individual RBD substitutions of B.1.617.1 (L452R, E484Q) and B.1.617.2/AY.1 (K417N, L452R, T478K) on D614G background. Titers against L452R (GMT 935 for Pfizer and 1781 for Moderna) and E484Q (GMT 798 for Pfizer and 1429 for Moderna) alone trended slightly lower. Likewise, titers against K417N (GMT 1208 for Pfizer and 2070 for Moderna) and T478K (GMT 1046 for Pfizer and 2113 for Moderna) alone or in L452R+T478K combination (GMT 964 for Pfizer and 1796 for Moderna) remained comparable to GMTs of WT(D614G) (Figure 2 and Supplementary Figure S1).

Similarly, GMT against B.1.1.7 (955 for Pfizer and 1917 for Moderna) and B.1.429 variant with the L452R substitution (1063 for Pfizer and 1799 for Moderna) was comparable to that of WT(D614G). Consistent with previous observations, B.1.351 variant (GMT 169 for Pfizer and 306 for Moderna) displayed ~7-fold lower titers compared to WT(D614G), whereas C.37, P.1, R.1 and B.1.526 variants displayed modestly reduced titers that are similar to the titers against B.1.617 pseudoviruses (GMT 452-707 for Pfizer and GMT 824-1332 for Moderna). Overall, the trends in neutralization titers for the vaccine-elicited sera against a large panel of variant pseudoviruses were similar to those for convalescent sera, though the GMTs were approximately 3 to 5-fold higher for vaccine-elicited sera.

A prior study showed that convalescent sera and vaccine-elicited antibody neutralization titers against B.1.617 pseudoviruses bearing L452R-E484Q-P681R spike substitutions were about 2-5-fold reduction [20]. In this study, depending upon the infecting variant, convalescent sera displayed a modest 2-4-fold reduction in neutralization titers for B.1.617.1 compared to WT(D614G), while the vaccine-elicited sera displayed only 2.1-fold reduction. Our findings using B.1.617.1 pseudoviruses with the full complement of spike substitutions rather than just the L452R-E484Q-P681R spike substitutions confirm and extend the prior findings [20]. Finally, the 1.8-3.4-fold reduction in neutralization in vaccination group against B.1.617.2 pseudoviruses seen in our study and others suggests that the two-dose mRNA vaccines could significantly contribute to protection against both Kappa and Delta variants [36,39].

### 3.3. Antigenic Cartography

We used antigenic cartography to compare the relative difference in neutralization of distinct variants, exploiting heterogeneity in individual antibody responses to identify antigenic differences among strains. Antigenic map dimensionality was tested in 1-5 dimensions using cross-validation. The 3D maps had only slightly greater predictive power compared to the 2D maps; both the 2D and 3D maps performed much better than 1D (Supplementary Table S1). The 2D antigenic maps are presented for convalescent infection sera (Figure 3A) and vaccine sera (Figure 3B). To evaluate reproducibility in positioning of antigens on the map, we created bootstrap confidence intervals in which n=10,000 antigenic maps were made by resampling sera with replacement (bottom panels in Figure 3A and 3B). Antigenic maps are also shown in 3D (Figure 3C and 3D). Maps made with convalescent sera from infection only by WT(D614G) or only by strains containing L452R mutation were not sufficiently robust for antigen positioning.

**Figure 3.**
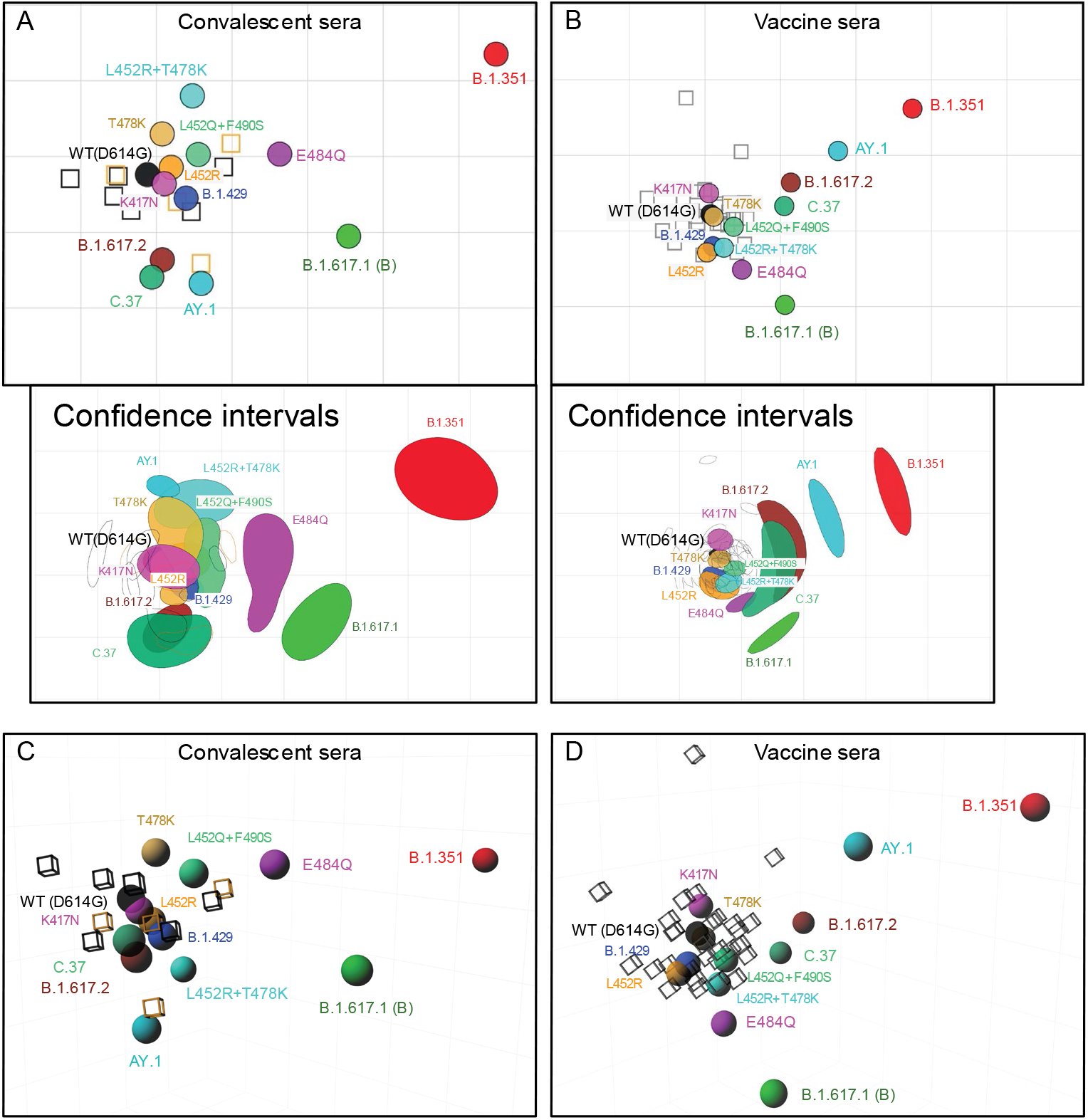
Antigenic maps of SARS-CoV-2 variants made using antigenic cartography. Antigenic maps were made separately of neutralizing antibody titers from (**A**, **C**) convalescent sera from individuals infected either WT(D614G) or strains with L452R mutation, or (**B**, **D**) sera from individuals vaccinated with either Pfizer/BioNtech BNT162b2 or Moderna mRNA-1273 vaccine. Sera are shown as open squares, pseudoviruses are shown as colored circles, labeled by strain name. Each grid-square corresponds to a two-fold dilution in the pseudovirus neutralization assay. Antigenic distance is interpretable in any direction. Antigenic maps shown with 70% bootstrap confidence intervals to convey uncertainty in positioning of antigens and sera were shown in the bottom panel of (**A**) and (**B**). Bootstrap confidence intervals were made with 10,000 resampled maps, each made by bootstrap sampling of antisera. (**A**) and (**B**): 2D map; (**C**) and (**D**): 3D map.

On the convalescent sera antigenic map, most of the pseudoviruses clustered close to the WT(D614G) pseudovirus and the WT(D614G) sera and L452R sera, including K417N, L452R, T478K, L452Q+F490S, and B.1.429 (0.25 to 0.72 antigenic units (AU) from WT (Table 3). B.1.617.2, C.37, and AY.1 clustered below WT(D614G) and slightly further away (1.14 – 1.60 AU), while L452R+T478K was also further away, but in the opposite direction (1.20 AU). However, in the bootstrap confidence interval 2D map and 3D map, L452R+T478K and AY.1 had distinct positions, clustering together with B.1.617.2 and C.37 in the 3D map and together but with T478K on the bootstrap 2D map. On all maps, B.1.351 was the most distinct from WT(D614G) (4.90 AU), followed by B.1.617.1 (2.78 AU), positioned between B.1.617.2 and B.1.351, and E484Q (1.77 AU), which was between WT(D614G) and B.1.351. These findings suggest that the full set of RBD substitutions in combination with substitutions outside the RBD contribute to antigenic difference of B.1.617.1, B.1.617.2, C.37, and B.1.351, while RBD substitutions alone in the WT(D614G) background do not reflect what is observed for each respective variant.

**Table 3.** Antigenic units (AU) from WT(614G) on 2D antigenic maps

| Pseudovirus | convalescent sera | Vaccine sera |
| --- | --- | --- |
| WT(D614G) | 0.00 | 0.00 |
| K417N | 0.25 | 0.28 |
| L452R | 0.33 | 0.51 |
| T478K | 0.57 | 0.05 |
| B.1.429 | 0.60 | 0.43 |
| L452Q-F490S | 0.72 | 0.35 |
| B.1.617.2 | 1.14 | 1.14 |
| L452R-T478K | 1.20 | 0.48 |
| C.37 | 1.35 | 0.98 |
| AY.1 | 1.60 | 1.89 |
| E484Q | 1.77 | 0.84 |
| B.1.617.1 | 2.78 | 1.55 |
| B.1.351 | 4.88 | 3.02 |

The antigenic map of the same SARS-CoV-2 variant pseudoviruses for vaccine-elicited sera was slightly different from the convalescent sera map. Again, pseudoviruses K417N, L452R, T478K, L452Q+F490S, and B.1.429 were close to WT(D614G) (0.05-0.51 AU). E484Q, C.37 and B.1.617.2 were further apart from WT(D614G) (0.84 to 1.14 AU) but were poorly coordinated and extended in elongated shapes around WT(D614G). B.1.617.1 was slightly more distant (1.55 AU) and was positioned adjacent to the other variants. Unexpectedly, AY.1 was between B.1.617.2 and C.37 and B.1.351 at 1.89 AU from WT(D614G), only slightly closer to WT(D614G) than B.1.351 was from WT(D614G) (3.02 AU). These antigenic maps reinforce what was observed in terms of neutralizing antibody titers. Vaccine-elicited sera noted a larger antigenic difference between AY.1 and WT(D614G) compared with convalescent sera and suggest AY.1 may have distinct antigenic properties compared with B.1.617.2. Notably, given that the vaccine-elicited sera had much higher titers than convalescent sera across variants, the titers against AY.1 were still higher in vaccinated individuals, meaning the antigenic difference may not translate into loss of vaccine protection. Furthermore, vaccine-elicited sera saw a smaller difference between B.1.351 and WT(D614G) than the convalescent sera, which may have aligned AY.1 and B.1.351 closer together. Overall, while these antigenic maps provide meaningful information on the relative positions of antigens, they are limited by the sera being so tightly clustered. Future antigenic maps with sera against distinct variants would enable more accurate evaluation of antigenic variation among the variants.

### 3.4. Spike RBD substitutions in B.1.617 variants affect sensitivity to therapeutic antibodies

We next evaluated 23 clinical-stage therapeutic neutralizing antibodies for potency against the B.1.617 variants. These antibodies were evaluated as part of the US Government COVID-19 response effort to inform the clinical testing and use of these antibodies [17]. Due to an agreement with the manufacturers who provided the antibodies, only blind codes are used to identify the antibodies. Amongst the therapeutic antibodies tested were thirteen monoclonal neutralizing anti-bodies (nAbs), six cocktail combinations (cnAbs) of two monoclonal anti-bodies and four polyclonal antibodies (pAbs).

B.1.617.1 pseudoviruses displayed complete resistance (>50-fold) to five nAbs (C, D, E, F and G) and partial resistance (10-50-fold) to one nAb (H) (Figure 4A and Supplementary Figure S2). The E484Q substitution alone conferred complete resistance (> 50 fold) to three nAbs (E, F and G), and partial resistance to one nAb (C). Several nAb classes were characterized previously to describe potential mechanisms of resistance. For instance, class 2 mAbs bind both up- and down-RBD configurations to prevent ACE2 binding as well as bridge adjacent down RBDs to lock spike in a closed prefusion conformation to further prevent S-ACE2 engagement [40]. E484K is a dominant resistance substitution for many RBD-directed nAbs including class 2 nAbs, potentially explaining resistance conferred by E484Q [40]. The L452R substitution conferred complete resistance to four nAbs (C, D, E and H). The L452R is centrally located in the receptor binding site and is a known resistance substitution for several monoclonal antibodies [24]. None of the cnAbs and pnAbs tested showed the loss of neutralization potency against L452R, E484Q, and B.1.617.1 (Supplementary Figure S2). Overall, E484Q and L452R substitutions fully accounted for the resistance of B.1.617.1 against these nAbs, and 17 of 23 tested therapeutic antibodies retained neutralization potency against B.1.617.1 (Supplementary Figure S2).

**Figure 4.**
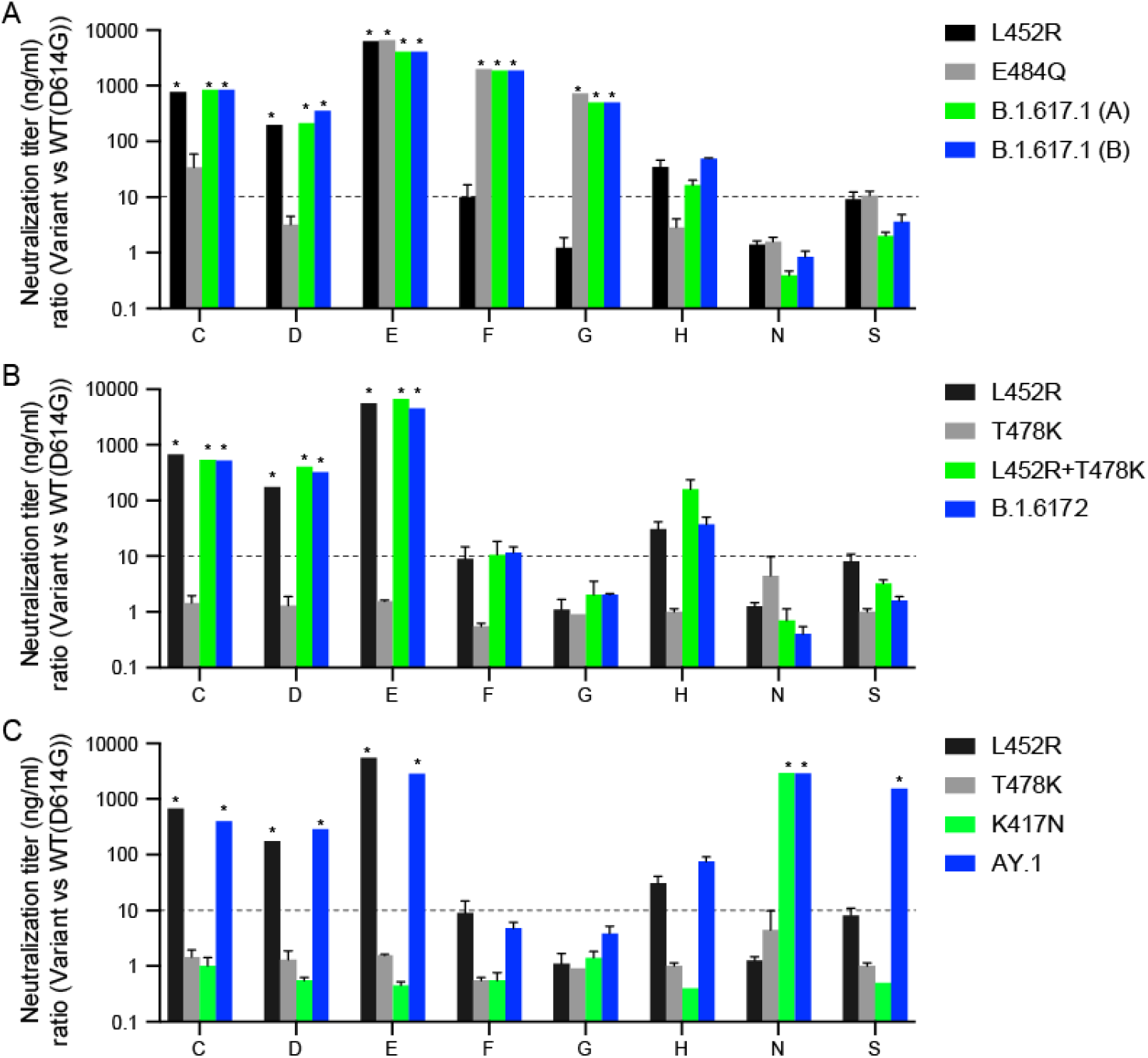
Neutralization of variant SARS-CoV-2 pseudoviruses by therapeutic antibodies. The antibody panel, consisting of 15 single neutralizing Abs (nAbs A-R), six combination Abs (S-X), and two polyclonal Abs (III-IV), were tested against SARS-CoV-2 pseudoviruses bearing the indicated spike proteins. Only those that displayed significant neutralization resistance compared to WT(D614G) are shown (Refer to Supplementary figure S2 for remaining antibodies). Antibodies are blinded according to an agreement with the manufacturers. The neutralization titer (IC_50_) ratios of the B.1.617.1 variants (**A**), B.1.617.2 variant (**B**), and AY.1 variant (**C**) relative to WT(D614G) are plotted. The dotted lines represent the neutralization titer ratio of 10. *: complete resistance at the highest concentration tested (>50-fold change of IC_50_).

B.1.617.2 pseudoviruses displayed complete resistance (>50-fold) to three nAbs (C, D, and E) and partial resistance (10-50-fold) to one nAb (H) (Figure 3B). The L452R substitution alone is responsible for the observed resistance of these nAbs (C, D, E and H) as pseudoviruses bearing L452R alone or in combination with T478K (L452R+T478K) displayed identical patterns of resistance as B.1.617.2 pseudoviruses. AY.1 pseudoviruses displayed similar resistance as B.1.617.2 above and also complete resistance to one additional nAb (N) due to the K417N RBD substitution and one cnAb (S) (Figure 4C).

### 3.5. B.1.617 pseudovirus infectivity and spike protein processing

Prior reports indicated increased infectivity of pseudoviruses containing the L452R substitution in spike in 293T-ACE2.TMPRSS2 cells due to L452R conferring enhanced RBD affinity to ACE2 [19,22]. We therefore investigated the impact of the B.1.617 spike substitutions on pseudovirus infectivity and found that B.1617.1, B.1.617.2, and AY.1 pseudoviruses, as well as pseudoviruses with single RBD L452R, T478K, and E484Q substitutions, displayed similar infectivity to WT(D614G) (Supplementary Figure S3).

Previous studies demonstrated that L452R and N501Y substitutions in RBD enhanced spike protein affinity to ACE2, which may have contributed to greater transmissibility of B.1.1.7, B.1.351, P.1 and B.1.427/429 variants [22,41,42]. To investigate whether the B.1.617 variants have an increased binding for ACE2, we measured the neutralization potency of human soluble ACE2 (sACE2) protein against the variant pseudoviruse. We found that the 50% inhibitory concentrations (IC_50_) of sACE2 against B.1.617.2 (IC_50_: 0.681 μg/ml; p < 0.01) was 4.2-fold lower compared to WT(D614G) (IC_50_: 2.88 μg/ml) (Table 4 and Supplementary Figure S4). In agreement with greater ACE2 affinity of L452R RBD compared to WT RBD as per SPR results reported previously, pseudoviruses containing the L452R substitution (IC_50_:1.189 μg/ml; p < 0.05) and L452R+T478K substitutions (IC_50_:1.228 μg/ml; p < 0.001) displayed 2.4- and 2.3-fold lower IC_50_. However, pseudoviruses with single spike RBD substitutions T478K or E484Q), as well as the B.1.617.1 (IC_50_:2.028 μg/ml) and AY.1 (IC_50_:1.966 μg/ml) spikes, displayed comparable IC_50_ to WT (D614G) (0.5-1.5-fold change) (Table 4 and Supplementary Figure S4). These findings suggest enhanced binding of Delta spike protein to ACE2 receptor, which may potentially contribute to greater infectivity and replication in ACE2-positive target cells.

**Table 4.** Neutralization of B.1.617 pseudoviruses by soluble ACE2

| Pseudovirus | IC<sub>50</sub> (µg/ml) | Fold change (vs WT(D614G)) | *P* value (vs WT(D614G)) |
| --- | --- | --- | --- |
| WT(D614G) | 2.880 | 1.0 |  |
| B.1.617.1 (B) | 2.028 | 1.4 | <0.01 |
| B.1.617.2 | 0.681 | 4.2 | <0.01 |
| AY.1 | 1.966 | 1.5 | n.s. |
| L452R | 1.189 | 2.4 | <0.05 |
| T478K | 4.266 | 0.7 | n.s. |
| L452R+T478K | 1.228 | 2.3 | <0.001 |
| E484Q | 5.568 | 0.5 | n.s. |

*P* values were calculated by One-way analysis of variance (ANOVA) with Dunnett's multiple comparisons tests (variants vs WT(D614G)). n.s.: not significant

Since P681H enhanced proteolytic processing of B.1.1.7 spike [26], we next evaluated spike proteolytic processing on B.1.617 variants carrying the P681R substitution. The B.1.617 variant pseudoviruses displayed efficient proteolytic processing of spike compared to Wuhan-Hu-1 (D614) and WT(D614G) as observed by the S1/S ratio (Figure 5A). We further evaluated the effect of P681R substitution on furin cleavage of SARS-CoV-2 by introducing this substitution in Wuhan-Hu-1 (D614) and WT(D614G) spike backgrounds. For all pseudoviruses bearing the P681R spike substitution, including B.1.617.1, B.1.617.2, Wuhan-Hu-1+P681R and P681R (+D614G), spike cleavage as determined by higher cleaved S1 subunit to total S (S1+S) ratios was enhanced compared to the respective original pseudoviruses (Figure 5A). B.1.1.7 and WT(D614G) pseudoviruses bearing P681H also displayed enhanced spike cleavage (Figure 5A), consistent with a previous report [26]. B.1.429 pseudoviruses that lack P681R or P681H substitutions near the furin cleavage site displayed inefficient spike cleavage.

**Figure 5.**
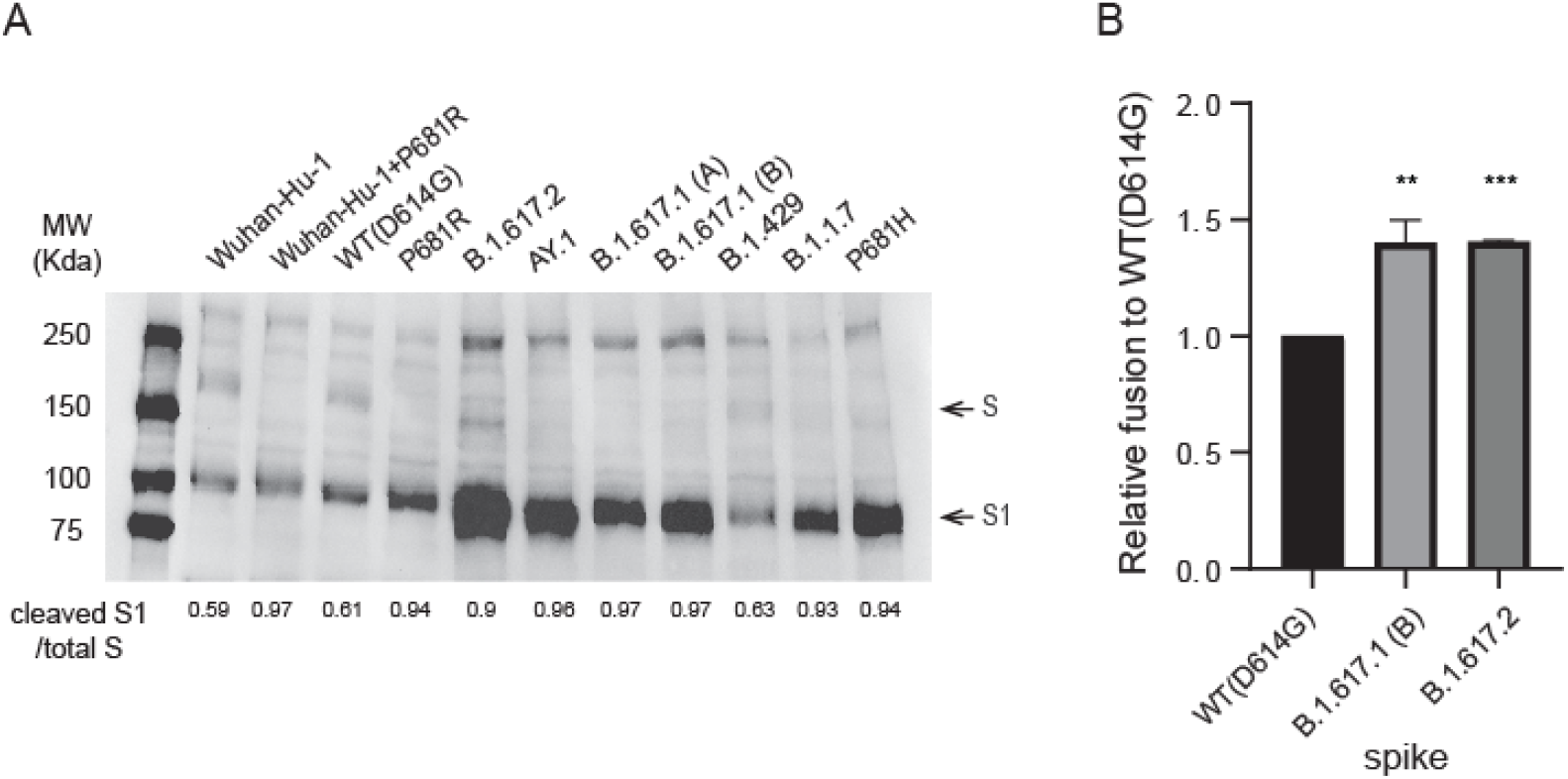
Proteolytic processing and fusogenic activity of B.1.617 variant spike proteins. (**A**) Western blot analysis of SARS-CoV-2 spike content of pseudoviruses. S/S1 was detected using a rabbit antibody against the SARS-CoV-2 S1 region. The image was a represent of three independent Western blots. (**B**) Spikes-induced cell-cell fusion quantified by β-galactosidase complementation assay. Y-axis indicates the relative β-galactosidase activity in variant spikes induced cell-cell fusion compared to WT(D614G) spike at 24 hour post co-culturing of spike-transfected 293T-ω cells and α-subunit-transfected 293T.ACE2.TMPRSS2 cells. X-axis indicates transfected spike in 293T-ω cells. Bars: Mean +/− SD of four independent experiments. P values were calculated by One-way analysis of variance (ANOVA) with Dunnett’s multiple comparisons tests (variants vs WT(D614G)). **: P≤0.01, ***: P≤0.001;

To gain further insight into the furin processing efficiency at the S1/S2 site of the B.1.617 S, we undertook a bioinformatic approach utilizing the PiTou and ProP furin cleavage prediction tools, comparing B.1.617 to the Wuhan-Hu-1 (D614) prototype spike and B.1.1.7 spike, as well as spike proteins of several lineage specific mammalian and animal CoVs. The PiTou algorithm combines a hidden Markov model and knowledge-based cumulative probability score functions for the functional characterization of a 20 amino acid cleavage motif from P14 to P6’ for furin binding and cleavage, whereas ProP predicts furin cleavage sites basing on experimental data-derived networks [43,44]. Both algorithms predicted a greater increase in the furin cleavage for B.1.617 lineage variants (PiTou: 12.4; Prop: 0.698) compared to Wuhan-Hu-1 (PiTou: 9.19; Prop: 0.62) and B.1.1.7 (PiTou: 9.9; Prop: 0.7) (Table 5) [44]. As expected, proteins not containing furin cleavage site displayed relatively lower score while much higher scores were shown for the proteins containing furin cleavage site [45,46].

**Table 5.**
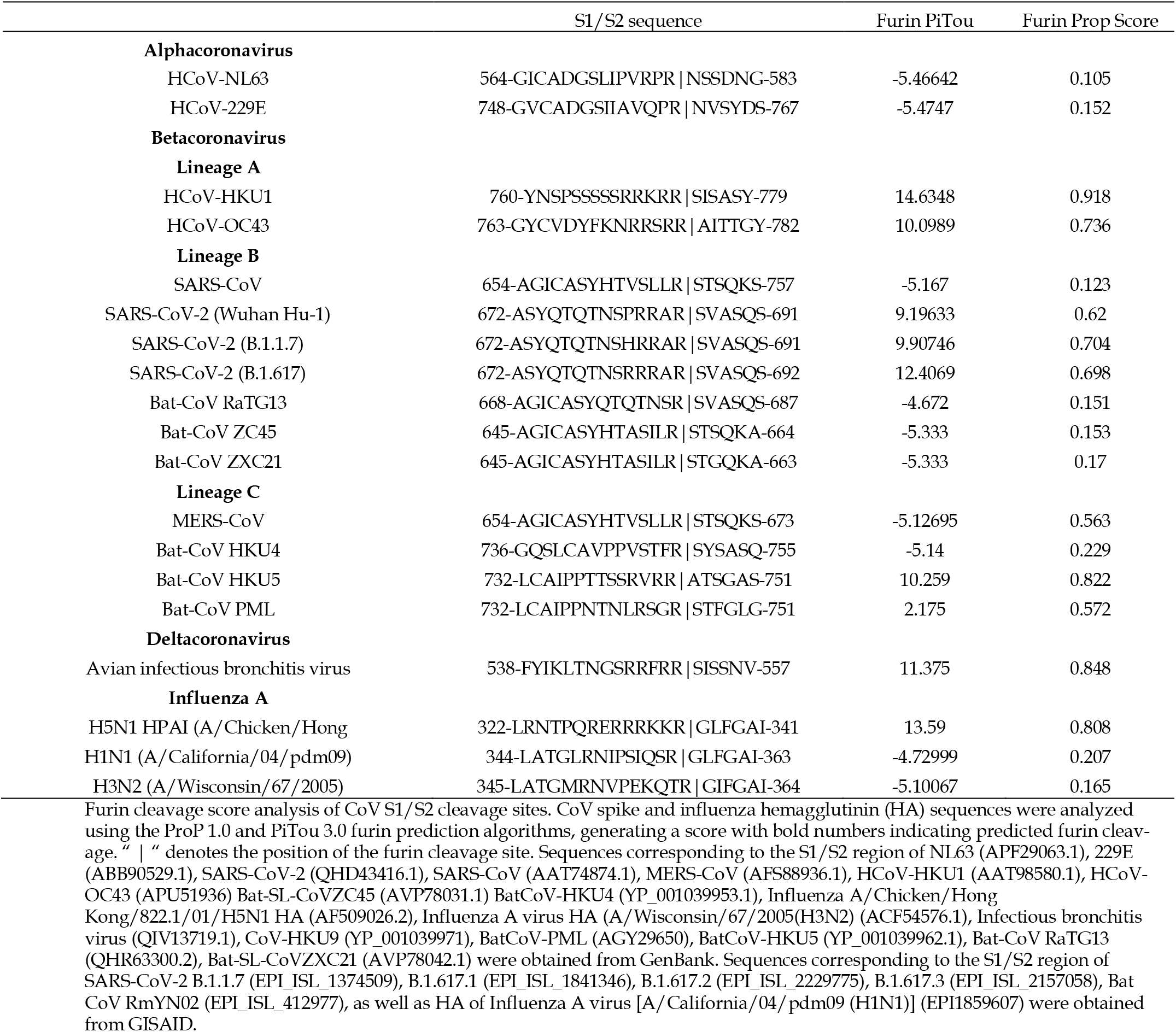
Prediction scores of furin cleavage at S1/S2 junction in coronavirus spikes

While SARS-CoV-2 S1/S2 P-R-R-A-R furin cleavage site conforms to a minimal furin recognition motif, R-X-X-R, the presence of H/R instead of P increases the total number of basic residues to four. This presence of basic residue H/R results in additional electrostatic and intramolecular hydrogen bonding to gain substrate turnover [47]. In fact, this basic residue is represented in several viral and cellular proteins [47,48]. While the cleavage is not necessary to enhance pseudovirus infectivity, the furin cleavage of spike has been shown to be necessary for the virus transmission as demonstrated in ferret model [49]. The P681H substitution in B.1.1.7 variants has been associated with enhanced virus transmissibility whereas the P681R substitution in B.1.617 variants has been associated with enhanced pathogenicity in a hamster model and transmissibility [26,49–51]. Further investigations of the role of P681H/R substitutions for virus transmissibility and cell-to-cell spread in *in vivo* animal models are needed. Furthermore, continued surveillance is necessary as several independent lineages have recently emerged containing additional substitutions proximal to S1/S2 cleavage junction, such as the B.1.2 and B.1.525 with Q677H, C.37 and B.1.617.2 with Q675H, and C.1.2 with N679K substitutions [52,53]. On the other hand, functions of furin cleavage site substitutions in host cell attachment and antigenicity cannot be excluded since amino acids N^679^- R^685^ and E^661^- R^685^ have been reported to have host neuropilin-1 attachment [54] and Staphylococcus enterotoxin B-like super-antigenic functions [55].

### 3.6. B.1.617 spike-mediated cell-cell fusion

To quantify whether higher ACE2 binding and furin cleavage of B.1.617.2 spike augments fusion between virus and/ or cell membranes, we performed cell-cell fusion assays by complementing β-galactosidase subunits in spike-transfected effector cells and 293T-ACE2.TMPRSS2 target cells. Compared to WT(D614G), both B.1.617.1 and B.1.617.2 spike induced significantly higher cell-cell fusion activity when controlled for spike cell surface expression (4000 MFI of spike protein on cell surface) (Figure 5B). Our findings extend previous studies indicating that the P681R substitution increases spike proteolytic cleavage and facilitates cell-cell fusion [54,55]. Our findings also suggest that fusogenic potential of spike proteins may be influenced by both ACE2 binding affinity as well as proteolytic cleavage of spike.

## 4. Conclusion

Here we show that pseudoviruses bearing B.1.617.1 spike with L452R and E484Q substitutions, and B.1.617.2 spike with K417N, L452R and T478K substitutions, have modestly reduced susceptibility to neutralization by Pfizer or Moderna vaccine-elicited sera and convalescent sera compared to pseudoviruses bearing WT(D614G) spike. The individual L452R, T478K, E484Q and dual L452R+T478K substitutions accounted for most but not all of the reduction in neutralization potency of the sera, suggesting contributions from substitutions in the NTD/CTD. Neutralization titers, as well as antigenic maps, indicated that the full set of RBD substitutions in combination with substitutions outside the RBD contribute to antigenic differences of B.1.617.1, B.1.617.2, and C.37 variants. Antigenic distances between the variants also tended to be more spaced apart in the map generated by the convalescent sera compared to the vaccine-elicited sera. Differences between convalescent and vaccine-elicited immune responses may be further impacted by the recent global dominance of B.1.617.2 and its sub lineages. Limitations in our study include the small numbers of sera samples in the convalescent and vaccine cohorts. Potential differences in COVID-19 severity in the convalescent sera cohort and time of sera collection could also affect neutralization titers. Nonetheless, most sera from convalesced and vaccinated individuals neutralized the B.1.617.1, B.1.617.2, and AY.1 variants. These findings with the vaccine-elicited sera suggest that the two-dose immunization with current mRNA vaccines will likely induce protective immunity against the tested B.1.617 variants. Furthermore, 17 of 23 therapeutic neutralizing antibodies retained complete neutralization against B.1.617 variants.

Resistance to the remaining therapeutic neutralizing antibodies is due to RBD substitutions, K417N, L452R, and E484Q, but not T478K. As B.1.617.2 variants continue to evolve, it will be important to continue to monitor how new substitutions in spike impact their resistance to therapeutic neutralizing antibodies and vaccine efficacy.

## Funding

This work was supported by the institutional funds from the US Food and Drug Administration and the Intramural Research Program of the National Institute of Allergy and Infectious Diseases.

## Informed Consent Statement

Informed consent was obtained from all subjects involved in the study.

## Acknowledgments

The authors would like to thank Hongquan Wan (FDA), and Hailun Ma (FDA) for the critical review of the manuscript.

## Conflicts of Interest

The authors declare no conflicts of interest.

## Notes

### Competing Interest Statement

The authors have declared no competing interest.

