## Supplemental Figures and Tables for "SARS-COV-2 Delta variant displays moderate resistance to neutralizing antibodies and spike protein properties of higher soluble ACE2 sensitivity, enhanced cleavage and fusogenic activity"

### Supplementary figures

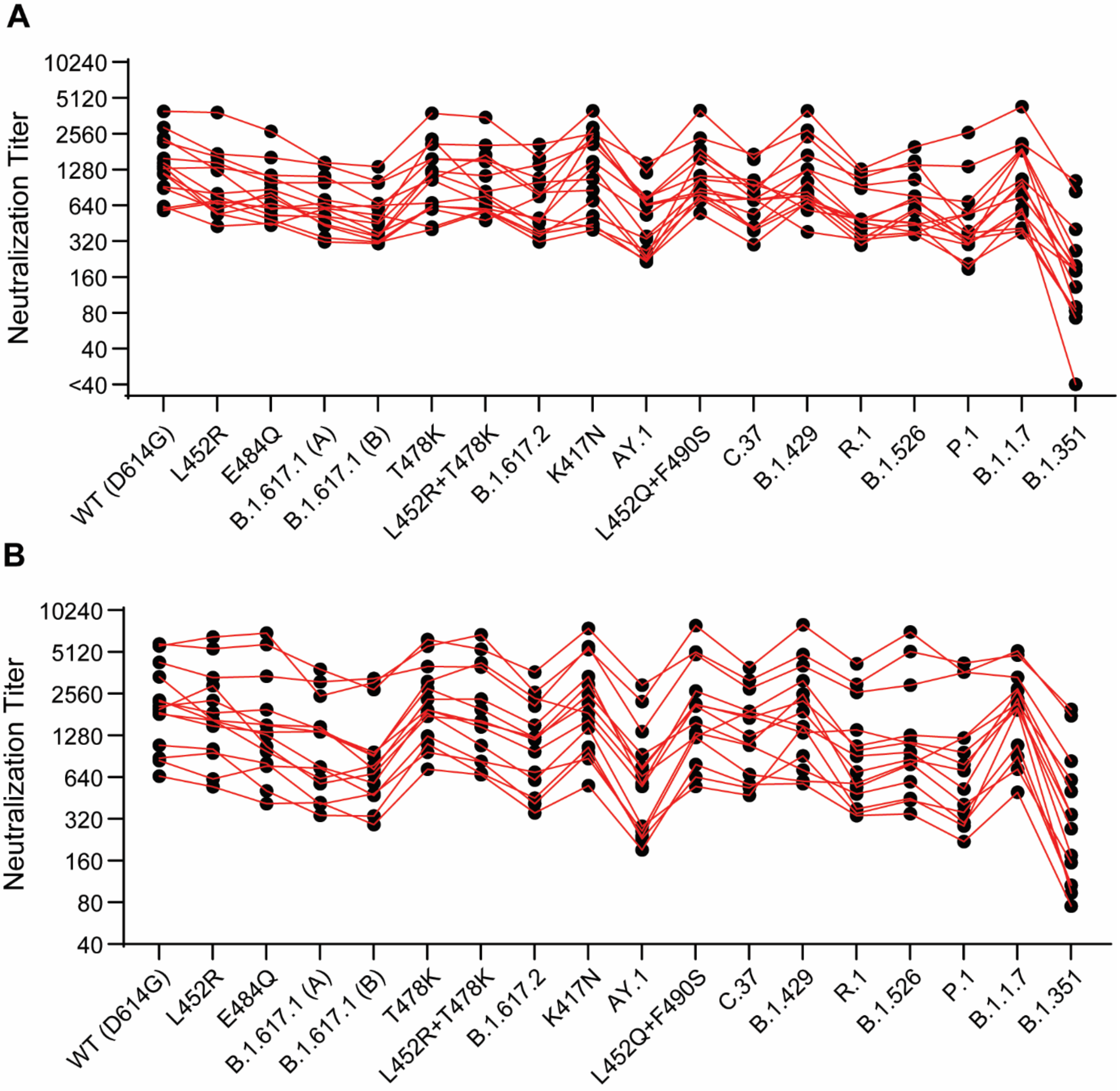


**Supplementary Figure S1. Neutralization of variant SARS-CoV-2 pseudoviruses by vaccination sera.** (**A**) Individual neutralization titers of Pfizer/BioNtech BNT162b2 vaccination sera are presented. (**B**) Individual neutralization titers of Moderna mRNA-1273 vaccination sera are presented. Lines connect responses from a given subject against the indicated variants.

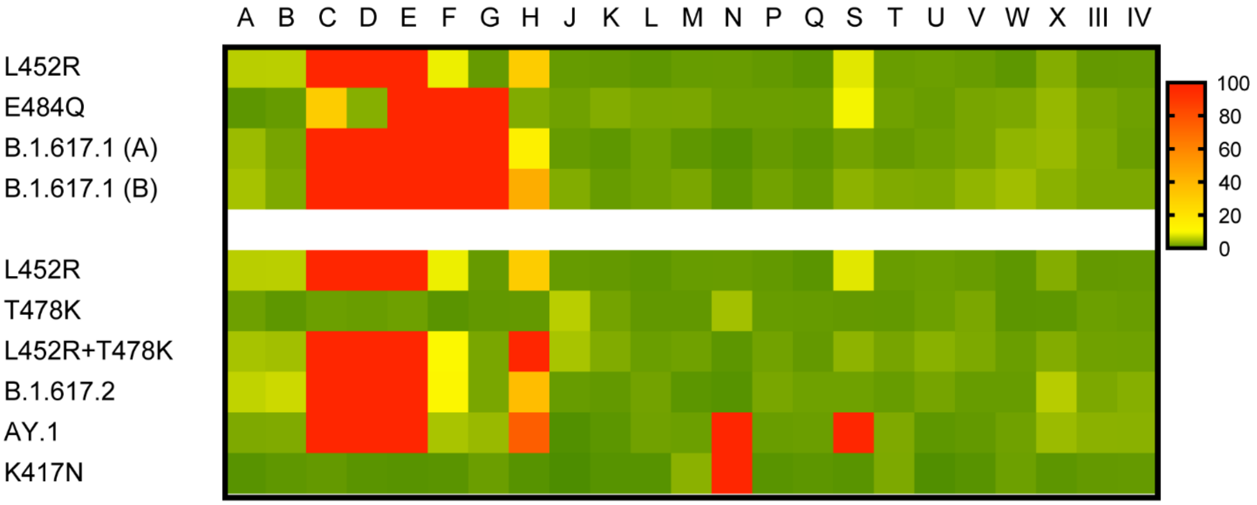


**Supplementary Figure S2. Neutralization of variant SARS-CoV-2 pseudoviruses by therapeutic antibodies.** The antibody panel, consisting of 15 single nAbs, 6 cnAbs, and 2 pAbs were tested against pseudoviruses bearing variant spikes and spikes with indicated RBD substitutions in the D614G background. Antibodies are blinded-coded according to an agreement with the manufacturers. Heat map representing the ratio of $IC_{50}$ values of the variant or RBD mutant spike relative to WT(D614G). Red indicates loss of potency ($IC_{50}$ ratios > 50). Yellow indicates moderate loss of potency ($IC_{50}$ ratios between 10-50), and green indicates retention of potency ($IC_{50}$ ratios <10).

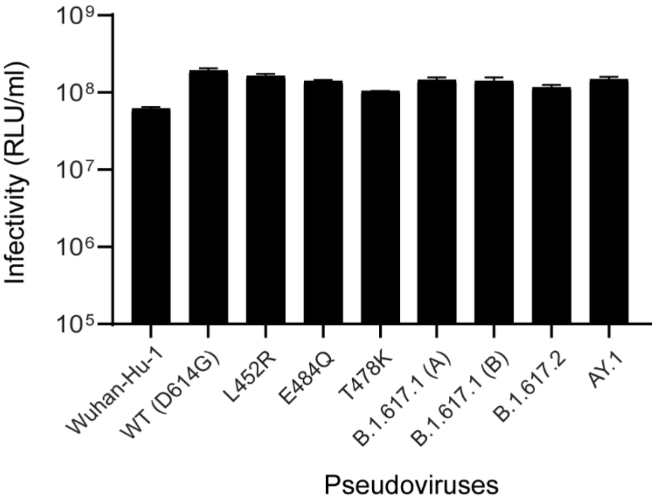


**Supplementary Figure S3. Infectivity of pseudoviruses with variants spikes.** Spike-bearing pseudoviruses infectivity in 293T-ACE2.TMPRSS2 cells. Cells were infected with pseudoviruses bearing spikes from SARS-CoV-2 variants. Bars: Mean +/- SD.

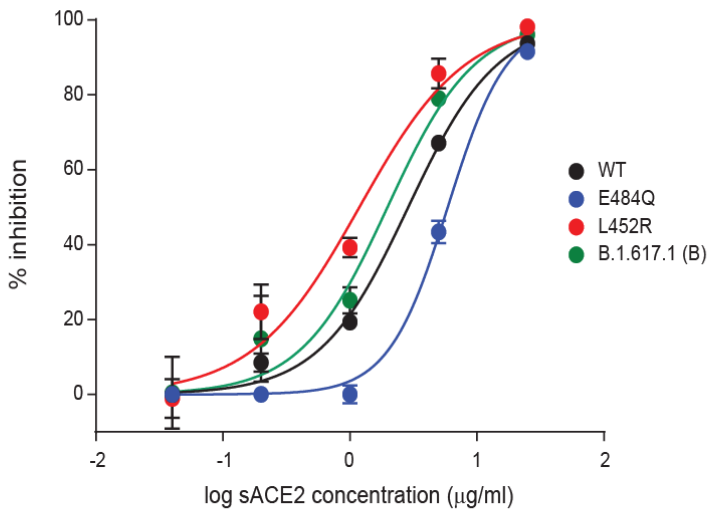


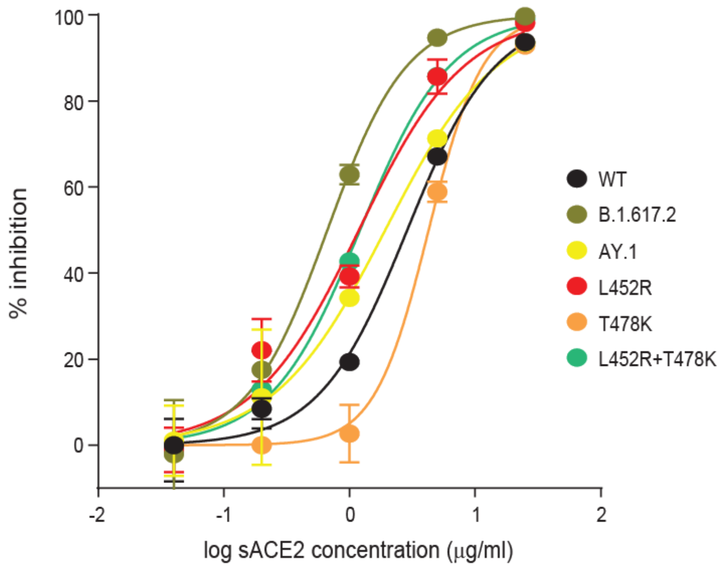


**Supplementary Figure S4. Neutralization of B.1.617 variant pseudoviruses by soluble ACE2.** Pseudoviruses bearing B.1.617.1 (**A**) and B.1.617.2 (**B**) variant spike proteins were incubated with a serially diluted soluble ACE2 (sACE2) and then applied to 293T-ACE2.TMPRSS2 cells for infection. Each curve indicates percent inhibition of pseudoviruses by sACE2. The image represents results of one of three independent experiments.

**Supplementary Table S1.** Dimensionality testing of Antigenic Cartography maps

| Dimension | Convalescent sera map | | | | Vaccine sera map | | | |
|---|---|---|---|---|---|---|---|---|
| | Measured titers | | Thresholded titers | | Measured titers | | Thresholded titers | |
| | Mean RMSE | variance RMSE | Mean RMSE | variance RMSE | Mean RMSE | variance RMSE | Mean RMSE | variance RMSE |
| 1D | 0.94 | 0.02 | 1.26 | 0.37 | 0.54 | 0.003 | 2.38 | 0.06 |
| 2D | 0.95 | 0.15 | 1.41 | 1.18 | 0.51 | 0.003 | 2.18 | 0.07 |
| 3D | 0.89 | 0.12 | 1.36 | 1.11 | 0.49 | 0.003 | 2.26 | 0.03 |
| 4D | 0.88 | 0.14 | 1.34 | 1.14 | 0.49 | 0.002 | 2.23 | 0.02 |
| 5D | 0.91 | 0.24 | 1.41 | 1.73 | 0.49 | 0.002 | 2.23 | 0.02 |

Dimensionality testing to identify best number of dimensions for fitting the antigenic maps. Each dataset was tested with cross-validation in 1-5 dimensions (100 maps, each made with 75% of the data to predict the exclude 25%), with low mean and variance in root-mean squared error (RMSE) for measured titers (within limit of detection) and thresholded titers (below the limit of detection) indicating the optimal number of dimensions. Convalescent sera from individuals infected either WT(D614G) or strains with L452R mutation. Vaccine sera from individuals vaccinated with either Pfizer/BioNtech BNT162b2 or Moderna mRNA-1273 vaccine.
